# Rapid specialization of counter defenses enables two-spotted spider mite to adapt to novel plant hosts

**DOI:** 10.1101/2020.02.26.966481

**Authors:** Golnaz Salehipourshirazi, Kristie Bruinsma, Huzefa Ratlamwala, Sameer Dixit, Vicent Arbona, Emilie Widemann, Maja Milojevic, Pengyu Jin, Nicolas Bensoussan, Aurelio Gómez-Cadenas, Vladimir Zhurov, Miodrag Grbic, Vojislava Grbic

## Abstract

Genetic adaptation, occurring over a long evolutionary time, enables host-specialized herbivores to develop novel resistance traits and to efficiently counteract the defenses of a narrow range of host plants. In contrast, physiological acclimation, leading to the suppression and/or detoxification of host defenses is hypothesized to enable broad-generalists to shift between plant hosts. However, the host adaptation mechanisms used by generalists composed of host-adapted populations are not known. *Tetranychus urticae* is an extreme generalist herbivore whose individual populations perform well only on a subset of potential hosts. We combined experimental evolution, *Arabidopsis* genetics, mite reverse genetics, and pharmacological approaches to examine mite host adaptation upon the shift of a bean-adapted population to *Arabidopsis thaliana*. We showed that cytochrome P450 monooxygenases are required for mite adaptation to *Arabidopsis*. We identified activities of two tiers of P450s: general xenobiotic-responsive P450s that have a limited contribution to mite adaptation to *Arabidopsis* and adaptation-associated P450s that efficiently counteract *Arabidopsis* defenses. In ≈25 generations of mite selection on *Arabidopsis* plants, mites evolved highly efficient detoxification-based adaptation, characteristic of specialist herbivores. This demonstrates that specialization to plant resistance traits can occur within the ecological timescale, enabling the two-spotted spider mite to shift to novel plant hosts.

## Introduction

During millions of years of co-evolution with their host plants, herbivores have developed two main strategies to counteract plant resistance traits. Specialist herbivores have evolved highly efficient adaptation mechanisms against a limited set of host defenses, including modified feeding behavior (Helmus and Dussourd, 2005), suppression of plant defenses (Zhao *et al*., 2015), reduced xenobiotic target site sensitivity (Dobler *et al*., 2012), sequestration (Beran *et al*., 2018), and detoxification of plant toxins (Ratzka *et al*., 2002). In contrast, generalist herbivores evolved an innate ability to feed on a broad range of host plants that display a wide array of resistance traits (Barrett and Heil, 2012; Despres *et al*., 2007; Heidel-Fischer and Vogel, 2015). The generalist lifestyle is considered to be ecologically and evolutionarily advantageous (Futuyma and Moreno, 1988; Loxdale and Harvey, 2016). However, the fitness trade-offs arising from the biochemical and physiological costs of polyphagy have been proposed to select for the specialization of herbivore–plant interactions (Dall and Cuthill, 1997). Generalist herbivores evolved two main strategies to counteract a diverse array of plant host defenses. Broad-generalists, where all individuals can feed on a wide range of plant hosts, rely on rapid transcriptional plasticity to colonise distantly related plants (Fenton *et al*., 1998; Mathers *et al*., 2017). However, the majority of generalist herbivores are regarded as composites of populations that themselves thrive on a subset of potential hosts (Barrett and Heil, 2012; Fox and Morrow, 1981; Peccoud *et al*., 2009). These populations exist in a wide range of host specializations and are proposed to ultimately lead to the formation of specialist species (Peccoud *et al*., 2009). Consistent with these theoretical predictions, specialist herbivores dramatically outnumber generalists (Rafter and Walter, 2020). Specialization to plant resistance traits is assumed to occur over a long evolutionary time and evokes complex adaptive genetic changes that result in the establishment of highly efficient adaptation mechanisms against a limited set of host defenses. Despite our understanding of host specialization in specialist herbivores, the mechanisms that mediate the formation of host-adapted populations in generalist herbivores are poorly understood.

Attenuation of plant responses induced by herbivore feeding has been proposed to be one of the mechanisms of host adaptation (Kant *et al*., 2008; Musser *et al*., 2012; Wu *et al*., 2012; Zarate *et al*., 2007). Consistently, effectors that modulate plant defenses have been identified in secretions of a number of generalist herbivores belonging to different feeding guilds (Bass *et al*., 2013; Basu *et al*., 2018; Hogenhout and Bos, 2011; Jonckheere *et al*., 2016; Kaloshian and Walling, 2016; Musser *et al*., 2002; Wu *et al*., 2012). Their mode of action is largely unknown but, to be effective against many host plants, it is assumed that they either target conserved compounds or pathways associated with plant defense or have a broad-spectrum activity with only a specific subset being effective against any particular host. Another general mechanism of host adaptation is metabolic resistance whereby herbivores effectively detoxify ingested plant toxins. In specialists, metabolic resistance is based on a limited number of detoxification enzymes that have high specificity and efficiency for a given plant toxin (Gloss *et al*., 2014; Heidel-Fischer *et al*., 2019; Li *et al*., 2003; Mao *et al*., 2006; *Ratzka et al., 2002*; Wittstock *et al*., 2004). Genes encoding these enzymes usually carry nucleotide substitutions in coding regions that increase their activity against the plant toxin (Dobler *et al*., 2012; Gloss *et al*., 2014; Wen *et al*., 2006) and/or in the promoter sequences resulting in their constitutive and high level of expression (Bass *et al*., 2013). In contrast, it is assumed that generalist herbivores rely on ubiquitous classes of detoxification enzymes (e.g. carboxyl/cholinesterases (CCEs), cytochrome P450 monooxygenases (CYP450s), glutathione S-transferases (GSTs), UDP-glycosyltransferases (UGTs), and ABC transporters (ABCs)) that were shown to accept structurally diverse substrates which they metabolize with low levels of activity (Halon *et al*., 2015; Li *et al*., 2004; Shi *et al*., 2018; Snoeck *et al*., 2019). Genes encoding general detoxification enzymes have undergone extensive amplification and neofunctionalization, and are transcriptionally responsive to a wide range of xenobiotics (Govind *et al*., 2010; *Grbic et al., 2011*; Muller *et al*., 2017; Schweizer *et al*., 2017; Wybouw *et al*., 2015; Zhurov *et al*., 2014). Thus, it is hypothesized that attenuation of plant defenses combined with the expanded and functionally versatile detoxification capabilities enable generalist herbivores to cope with diverse allelochemicals and to feed on many host plants.

The two-spotted spider mite (TSSM) *Tetranychus urticae* is an extreme generalist herbivore that feeds on over 1,100 plant species from more than 100 families (Migeon and Dorkeld, 2021). Such a wide host range indicates that TSSM can counteract a great diversity of plant resistance traits. However, individual TSSM populations do not perform equally well on all potential host plants (Agrawal *et al*., 2002; Fellous *et al*., 2014; Fry, 1989). Instead, TSSM has an outstanding ability to adapt to new hosts (Fry, 1989; Gould, 1979; Magalhaes *et al*., 2007; Wybouw *et al*., 2015). The mechanism of this host adaptability is not known. The analysis of plant transcriptional changes following the host shift revealed that some TSSM populations can suppress plant induced responses (Kant *et al*., 2008; Wybouw *et al*., 2015). At the same time, TSSM massively reprograms its detoxification capacity (Dermauw *et al*., 2013; Wybouw *et al*., 2015; Zhurov *et al*., 2014) and the complement of its salivary secretions (Jonckheere *et al*., 2018; Jonckheere *et al*., 2016; Villarroel *et al*., 2016). However, no functional evidence explains whether these changes contribute to TSSM host adaptation or if they merely reflect stress responses or different feeding physiology due to the host shift. In addition, it remains unclear if changes to the initial xenobiotic responses are sufficient for TSSM adaptation to a new host, or if additional changes are required for the evolution of TSSM host adaptation.

Even though plants belonging to the Brassicaceae family have been recorded as TSSM plant hosts (Migeon and Dorkeld, 2021), *Arabidopsis thaliana* is a non-preferred host for the TSSM London reference population that is reared on a bean – *Phaseolus vulgaris* (Zhurov *et al*., 2014). In an experimental evolutionary setup, we adapted the TSSM London ancestral population to *Arabidopsis*. Using functional pharmacological and reverse genetics experiments that independently suppressed the activities of families of mite detoxification enzymes we provide, for the first time, *in vivo* functional evidence that *T. urticae* requires cytochrome P450 monooxygenase activity for adaptation to *Arabidopsis*. We identified activities of two tiers of P450s: general xenobiotic-responsive P450s that have a limited contribution to mite adaptation to *Arabidopsis* and adaptation-associated P450s that efficiently counteract *Arabidopsis* defenses. Our data reveal that in ≈25 generations, mites established a highly efficient detoxification response to *Arabidopsis* defenses, demonstrating that specialization to plant resistance traits can occur within an ecological timescale.

## Results

### TSSM feeding induces complex JA-regulated *Arabidopsis* defenses

To understand the mechanism of TSSM adaptation to *Arabidopsis*, we first characterized the complexity of *Arabidopsis* defenses that mites have to overcome in order to use *Arabidopsis* as a host. Jasmonic acid (JA) and its bioactive conjugate jasmonoyl-isoleucine (JA-Ile), Figure 1A, were shown to be required for *Arabidopsis* defenses against TSSM herbivory (Zhurov *et al*., 2014). A dramatic 5-fold decrease of mite fecundity on methyl jasmonate (MeJA) treated Col-0 plants, Figure 1B, indicates that JA-induced defenses are sufficient for *Arabidopsis* resistance against TSSM. JA responsiveness is mediated by the MYC2, MYC3, and MYC4 (MYC2,3,4) transcriptional activators (Devoto *et al*., 2005; Fernandez-Calvo *et al*., 2011), and consistently, *myc2 myc3 myc4* (*myc2,3,4*) mutant plants are extremely susceptible to TSSM, Figure 1B. Among MYC2,3,4-regulated genes are *CYP79B2* and *CYP79B3* that are required for the synthesis of Trp-derived secondary metabolites (Figure 1A, (Hull *et al*., 2000; Mikkelsen *et al*., 2000; Schweizer *et al*., 2013). One class of these metabolites are indole glucosinolates, shown to protect *Arabidopsis* plants against herbivory of a wide range of arthropods, including mites (Elbaz *et al*., 2012; Hopkins *et al*., 2009; Kim and Jander, 2007; Kim *et al*., 2008; Whiteman *et al*., 2012; Whiteman *et al*., 2011; Widemann *et al*., 2021; Zhurov *et al*., 2014). The modest increase in fecundity when TSSM fed on *cyp79b2 cyp79b3* (*cyp79b2,b3)* plants, Figure 1B, indicates that besides indole glucosinolates, there are other MYC2,3,4-regulated defenses that prominently protect *Arabidopsis* plants against mites. Thus, there are at least two distinct classes of JA-regulated *Arabidopsis* resistance traits against TSSM that are *CYP79B2 CYP79B3-*dependent and *-* independent.

**Figure 1.**
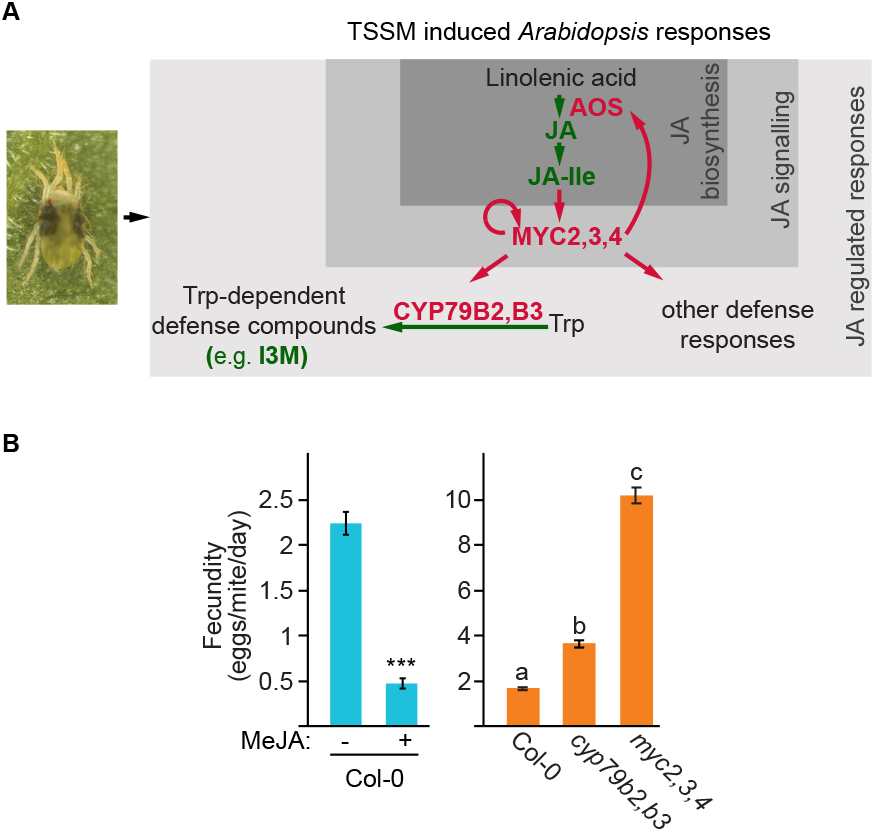
Two-spotted spider mite (TSSM) *Tetranychus urticae* induces complex defense responses in *Arabidopsis*. **A**, A simplified schematic of TSSM feeding-induced *Arabidopsis* defense responses. **B**, The fecundity of bean-reared TSSM (ancestral population) upon feeding on MeJA treated Col-0 plants (left) and on Col-0 wild type, *cyp79b2 cyp79b3* and *myc2 myc3 myc4* mutant plants *(myc2,3,4)* (right). Data are presented as mean number of eggs laid by a female mite per day ± SEM. Left, *n* = 18; asterisk indicates a significant difference between treated and control samples (ANOVA: ***P < 0.001). Right, *n* = 15 Different letters represent significant difference between means (Tukey’s HSD test, α = 0.05).

### Mites can adapt to a complex array of *Arabidopsis* defenses

To test if TSSM can adapt to the complex array of *Arabidopsis* defenses, we used an experimental evolutionary setup where an ancestral TSSM population that is highly susceptible to *Arabidopsis* defenses (Zhurov *et al*., 2014) was transferred and continuously maintained on *Arabidopsis* plants. The improvement of mite fitness on a new host has been reported to occur already after 5-10 generations (Agrawal, 2000; Fry, 1989; Fry, 1990), however, the improvement is gradual and reaches a stable state after 15-20 generations, depending on the ancestral population and the plant host (Agrawal, 2000; Fry, 1989; Magalhaes *et al*., 2009). We have initiated the analysis of mite adaptation after 18 months (≥25 generations) of selection, Figure 2A, to ensure consistency of mite adaptation state throughout the analysis. We initially infested *Arabidopsis* plants with approximately 1,000 fertilized female mites in a triplicated experiment on two *Arabidopsis* genotypes: a) Col-0, fully defended, wild-type plants, and b) *cyp79b2 cyp79b3* mutant plants that lack Trp-derived compounds but display the remaining JA-regulated defenses, Figures 1A and 2A. The performance of the ancestral and selected mite populations was subsequently quantified by counting the total number of eggs, larvae, nymphs, and adults derived from 20 adult females in 7 days. The performance was measured in two experimental regimes: a) direct transfer, where the initial mites were moved from their corresponding rearing plant hosts directly to the experimental plants, and b) indirect transfer, which included mite maintenance on bean plants for two generations before their transfer to the experimental plants. The performance of the ancestral and selected mite populations on bean, or *Arabidopsis cyp79b2 cyp79b3* and Col-0 plants had similar patterns in both direct (Supplemental Figure S1) and indirect (Figure 2B) transfer regimes, indicative of genetic adaptation that is independent of maternal and environmental physiological effects.

**Figure 2.**
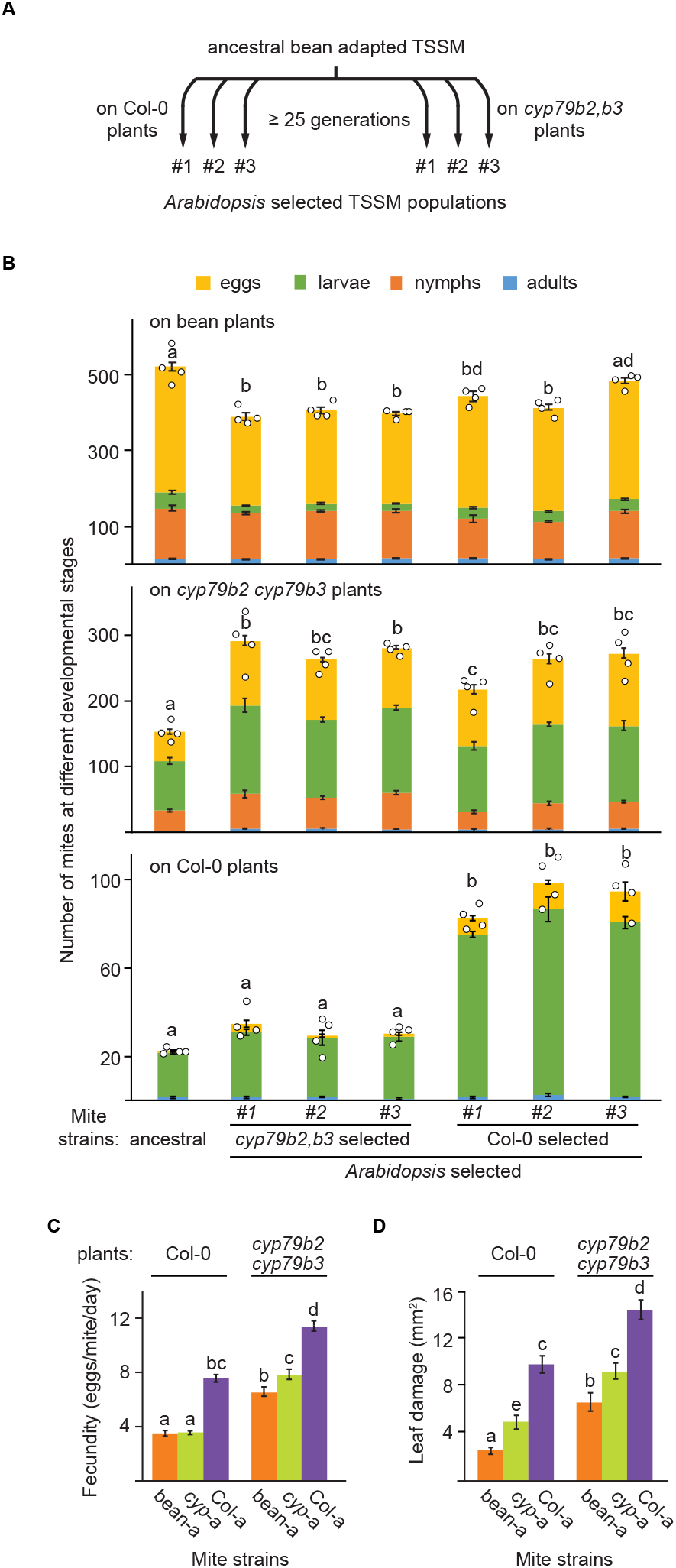
*Tetranychus urticae* can adapt to complex array of *Arabidopsis* induced defenses. **A**, A schematic of the experimental evolution procedure. **B**, Performance of ancestral and selected populations on bean (top), *cyp79b2 cyp79b3* (middle) and Col-0 (bottom) plants. Data are from the indirect transfer experiment where *Arabidopsis* selected TSSM were reared for two generations on bean before transfer to the experimental plants. Data represent the mean size of population (the sum of eggs, larvae, nymphs, and adults) derived from twenty adult female mites at seven days post infestation ± SEM, *n* = 4. Different letters represent significant difference between means (Tukey’s HSD test, α = 0.05). Data from direct transfer experiments where *Arabidopsis* selected TSSM were directly transferred from their rearing to the experimental plants is shown in Supplemental Figure S1. **C**, Fecundity of the ancestral (bean-a) and populations #3 of *Arabidopsis*-selected mites (*cyp*-a and Col-a) upon feeding on Col-0 and *cyp79b2 cyp79b3* plants. Data are presented as mean number of eggs laid by a female mite per day ± SEM, *n* = 30. **D**, Leaf damage of Col-0 and *cyp79b2 cyp79b3 Arabidopsis* plants upon herbivory of ancestral (bean-a), and populations #3 of *Arabidopsis*-selected mites (*cyp*-a and Col-a). Data represent mean area of chlorotic spots (mm^2^) ± SEM, *n* = 18. Different letters represent significant differences between means (Tukey’s HSD test, α = 0.05). Individual sample values are shown as open circles for *n* ≤ 10.

On bean plants, Figure 2B top, *Arabidopsis*-selected mite populations had slightly but significantly lower performance relative to the ancestral population, indicating that TSSMs can evolve the ability to exploit different hosts without a major reduction in their performance on the ancestral plants. On *cyp79b2 cyp79b3* plants, Figure 2B middle, the two *Arabidopsis*-selected mite populations performed significantly better than the ancestral population, suggesting that they can overcome *CYP79B2 CYP79B3*-independent *Arabidopsis* defenses to a similar extent. On Col-0 plants, Figure 2B bottom, only mites selected on Col-0 had increased performance over the ancestral population, showing that Col-0 selected mites were also able to adapt to *CYP79B2 CYP79B3-* dependent defenses. However, mites selected on *cyp79b2 cyp79b3* plants and the ancestral mite population that were not exposed to *CYP79B2 CYP79B3*-dependent *Arabidopsis* defenses were susceptible to these defenses and had similar and low performances when they fed on Col-0 plants.

We have arbitrarily chosen populations #3 of *cyp79b2 cyp79b3* and Col-0 selected mites for subsequent studies that were all performed in indirect transfer regimes. The analysis of additional mite fitness parameters, fecundity, and *Arabidopsis* leaf damage caused by mite feeding (Figure 2C and D, respectively) confirmed the adaptation status of these mite populations (from now on referred to as *cyp*-a (for #3 *cyp79b2 cyp79b3* selected mite population), Col-a (for #3 Col-0 selected mite population) and bean-a (for the ancestral bean-adapted London TSSM population). Thus, our data demonstrate that TSSM can adapt to novel plant hosts that have a complex array of defenses. Importantly, *Arabidopsis*-adapted mite populations retained high fitness on bean plants, thus, they expanded their host range without losing the ability to feed on the ancestral host. Moreover, mite adaptation to *CYP79B2 CYP79B3*-independent *Arabidopsis* defenses (present in both types of *Arabidopsis*-selected mite populations) can be uncoupled from mite adaptation to Trp-derived defenses (present only in mites selected on Col-0 plants).

### Ancestral and *Arabidopsis*-adapted mite strains induce similar JA-regulated *Arabidopsis* defenses

Suppression of induced plant defenses was proposed to be one of the mechanisms of TSSM adaptation to new host plants (Blaazer *et al*., 2018; Jonckheere *et al*., 2018; Jonckheere *et al*., 2016; Villarroel *et al*., 2016). One of the hallmarks of defense-suppressing TSSM populations is the similarity of their performance on wild-type and defense-deficient host plants (Kant *et al*., 2008). However, *Arabidopsis*-adapted mites, like the ancestral population, performed significantly better on *cyp79b2 cyp79b3* than on Col-0 plants, Figure 2B and C, suggesting that mite adaptation to *Arabidopsis* is not based on the suppression of host defenses. To corroborate the lack of defense suppression in *Arabidopsis*-adapted mites we determined the expression levels of JA-responsive *AOS, MYC2, CYP79B2*, and *CYP79B3* genes (labeled in red in Figure 1A) and the abundance of JA, JA-Ile, and indol-3-ylmethylglucosinolate (I3M) (labeled in green in Figure 1A) that were previously established as reliable markers of induced JA-regulated *Arabidopsis* defenses against mite herbivory (Widemann *et al*., 2021; Zhurov *et al*., 2014). The expression levels and the abundance of defense-associated markers were determined in Col-0 plants that were challenged with bean-a, *cyp*-a, or Col-a mites after 24 h of mite herbivory. As seen in Figure 3A, *cyp*-a and Col-a mites induced all JA-regulated marker genes to similar or higher levels relative to non-adapted bean-a mites. Likewise, JA, JA-Ile, and I3M accumulated at comparable levels in bean-a, *cyp*-a, and Col-a challenged Col-0 plants (Figure 3B). Even though we can not rule out the possibility that some plant pathways with minor effects on mite fitness may have been suppressed, our data suggest that ancestral and *Arabidopsis*-adapted mites are exposed to similar JA-regulated *Arabidopsis* defenses, shown to be both necessary and sufficient to protect *Arabidopsis* plants against mite herbivory.

**Figure 3.**
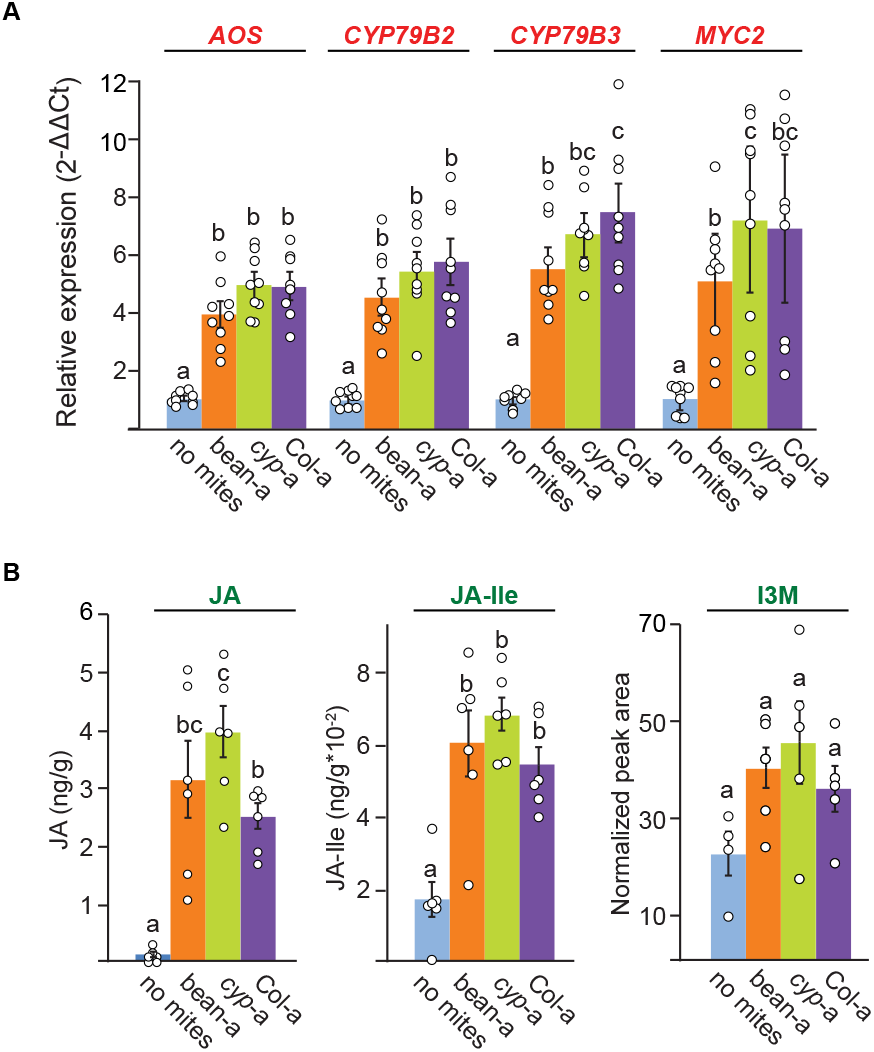
Bean-a, *cyp*-a and Col-a mites induce similar *Arabidopsis* defense responses. **A**, Expression of *AOS, CYP79B2, CYP79B3* and *MYC2* genes in Col-0 leaves in response to bean-a, *cyp*-a and Col-a mite feeding. Shown are means ± SEM of fold changes of expression levels detected by RT-qPCR relative to no mite control, *n* = 9. Primer sequences and amplification efficiencies (E) used in qPCR are shown in Supplemental Table 1. *PEROXIN4* (*AT5G25760*), was used as the reference gene. **B**, Levels of JA and JA-Ile (ng g^−1^ fresh weight), and relative level of the indol-3-ylmethyl glucosinolate (I3M, shown as normalized peak area), in 3-week-old Col-0 plants after herbivory of bean-a, *cyp*-a and Col-a mites for 24 h. Values are means ± SEM, *n* = 6. Different letters represent significant differences between means (Tukey’s HSD test, α = 0.05). Individual sample values are shown as open circles for *n* ≤ 10.

### Initial responses to *Arabidopsis* xenobiotics are similar in ancestral and *Arabidopsis*-adapted mites

The constitutive overexpression and/or increased transcriptional plasticity of genes encoding xenobiotic-metabolizing enzymes have been previously implicated in the evolution of TSSM resistance to pesticides (Kwon *et al*., 2015; Van Leeuwen and Dermauw, 2016; Van Leeuwen *et al*., 2010) and mite adaptation to several plant hosts (Agrawal *et al*., 2002; Wybouw *et al*., 2012; Wybouw *et al*., 2014; Wybouw *et al*., 2015). We have previously identified forty genes, primarily encoding cytochrome P450 monooxygenases (P450s) and UDP-glycosyltransferase (UGTs), that were associated with the initial TSSM responses to varying levels of Trp-derived defense compounds (Zhurov *et al*., 2014). To test if these genes may have contributed to mite adaptation to *Arabidopsis*, we have arbitrarily chosen three genes within each class *(CYP392A1, CYP392A16, CYP392D8, UGT201A2v2, UGT204B1*, and *UGT204A5)* and have used RT-qPCR to determine their expression in bean-a, *cyp*-a, and Col-a mites that were kept on bean plants for two generations before they were transferred to bean and *Arabidopsis* plants with (Col-0) or without (*cyp79b2 cyp79b3*) Trp-derived defense compounds. On the bean plants, all marker genes had similar and low expression levels in both bean-a and *Arabidopsis*-adapted mites, Figure 4, indicating that none of the tested genes underwent constitutive upregulation during mite adaptation to *Arabidopsis*. Consistent with the previous report (Zhurov *et al*., 2014), the transfer of bean-a mites to *cyp79b2 cyp79b3* or Col-0 plants resulted in induced expression of all tested genes. Their expression was also induced in *cyp*-a and Col-a mites when they were shifted from bean to *Arabidopsis* plants. Even though there was variability in the relative expression of chosen *CYP* and *UGT* genes in these mites, their levels were either comparable or lower than in bean-a mites (Figure 4). Thus, neither higher constitutive expression nor greater transcriptional plasticity of tested *CYP* and *UGT* genes associates with the *Arabidopsis*-adapted mites, demonstrating that: a) *Arabidopsis*-adapted mites retained responsiveness to *Arabidopsis* xenobiotics, and b) the expression of tested mite genes does not associate with TSSM adaptation to *Arabidopsis;* instead, it associates with general xenobiotic responsiveness of bean-a, *cyp*-a, and Col-a mites to shift from bean to *Arabidopsis* plants.

**Figure 4.**
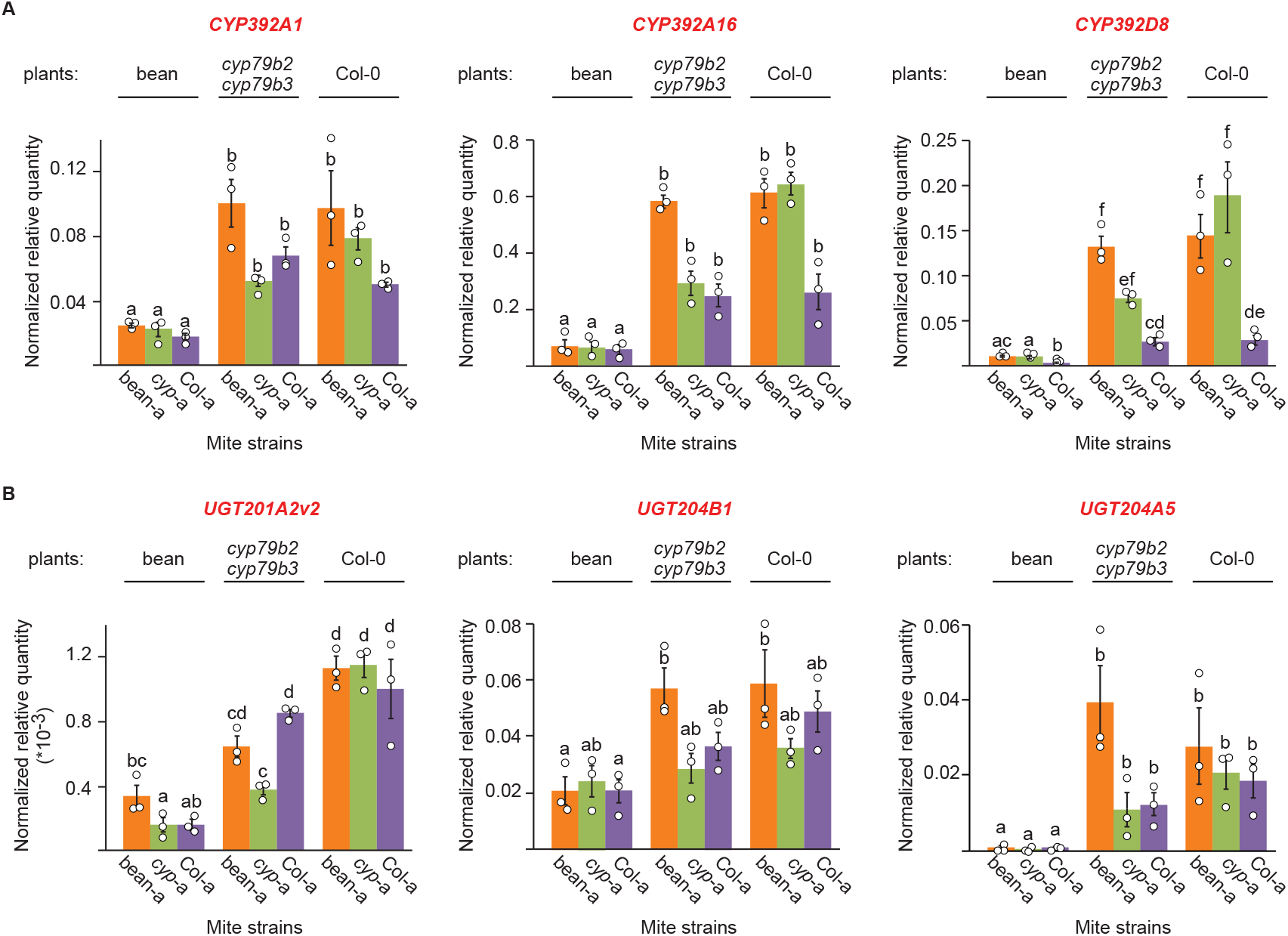
Levels of *Arabidopsis*-induced mite *cytochrome P450* (*CYP*) and *UDP-glycosyltransferase* (*UGT*) genes. **A**, *CYP392A1* (*tetur07g06410*), *CYP392A16* (*tetur06g04520*) and *CYP392D8* (*tetur03g05070*). **B**, *UGT201A2v2* (*tetur02g02480*), *UGT204B1* (*tetur02g09830*) and *UGT204A5* (*tetur05g00090*). Shown are means ± SEM for relative quantity of expression normalized to *RP49* (*tetur18g03590*) reference gene, *n* =3. Primer sequences and amplification efficiencies (E) used in qPCR are shown in Supplemental Table 1. Different letters represent significant differences between means (Tukey’s HSD test, α = 0.05). Individual sample values are shown as open circles for *n* ≤ 10.

### Metabolic resistance underlies TSSM adaptation to *Arabidopsis*

To identify changes in the detoxification potential associated with mite adaptation to *Arabidopsis*, we determined global enzymatic activity for three main protein families - esterases, glutathione-S-transferases (GSTs), and P450s - in ancestral and *Arabidopsis*-adapted mite populations feeding on bean and *Arabidopsis* plants. Overall, activities of all three enzymatic classes were responsive to the host plant challenge: they were lowest when mites fed on bean plants, and they progressively increased with the complexity of *Arabidopsis* defenses, Supplemental Figure S2. However, of the three classes of detoxification enzymes, P450 activity was consistently higher in Col-a mites across all plant hosts, indicating that both constitutive and inducible levels of P450 activity increased in Col-a mites, (three-way ANOVA; plant host : mite strain interaction F = 103.93, p = 5.451e-16, followed by Tukey’s HSD test, p < 0.01, n = 4).

To test the requirement of esterase, GST, and P450 activities for TSSM adaptation to *Arabidopsis*, we used S,S,S tributyl-phosphorotrithioate (DEF, an inhibitor of esterase activity), diethyl maleate (DEM, an inhibitor of GST activity), and piperonyl butoxide (PBO) and trichlorophenylpropynyl ether (TCPPE) (inhibitors of P450 activity) to reduce the activity of the indicated enzymatic classes. If a particular class of enzymes is required for mite adaptation to *Arabidopsis*, then the inhibition of its activity is expected to restore the susceptibility of *Arabidopsis*-adapted mites to *Arabidopsis* defense compounds. Concentrations of 2000 mg/L (DEM), 100 mg/L (DEF), 1000 mg/L (PBO), and 1500 mg/L (TCPPE) were used as they cause <10% mite mortality (Supplemental Figure S3), but were nevertheless capable of significantly reducing the corresponding enzymatic activities in Col-a mites feeding on Col-0 plants (Figure 5A,C and Supplemental Figure S4). As inhibitors do not affect all enzymes equally within the targeted enzymatic class (Feyereisen, 2014), the lack of inhibitory effect on mite performance is not strong evidence against the involvement of a particular enzymatic class in mite host adaptation. Conversely, the significant decrease of mite performance on a new host plant upon the application of a given inhibitor strongly supports the requirement of enzyme(s) within the corresponding class for mite host adaptation.

**Figure 5.**
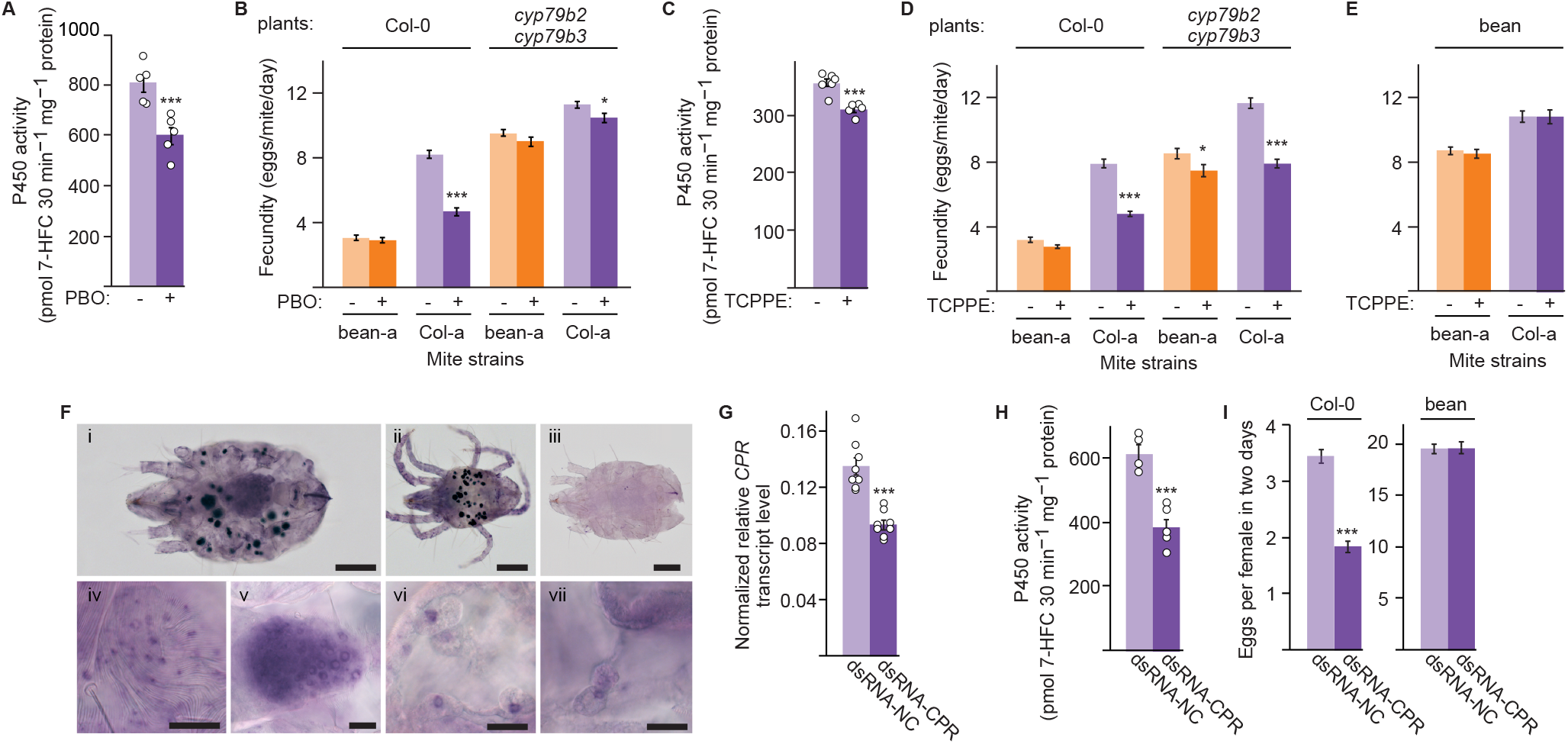
Cytochrome P450 activity is required for *Tetranychus urticae* adaptation to *Arabidopsis*. **A,C** The activities of cytochromes P450 in Col-a mites feeding on Col-0 plants after the application of piperonyl butoxide (PBO, in **A**) and trichlorophenylpropynyl ether (TCPPE, in **C**) inhibitors of P450 activity. Data are represented as mean ± SEM, *n* = 5. **B**,**D** The effects of PBO (**B**) and TCPPE (**D**) on fecundity of bean-a and Col-a mites upon feeding on Col-0 and *cyp79b2 cyp79b3* plants. **E**, The effect of TCPPE treatment on fecundity of bean-a and Col-a mites upon feeding on bean plants. Fecundity in **B**,**D** and **E** was measured over six days and data are presented as mean number of eggs laid by a female mite per day ± SEM; *n* = 30. **F**, Whole-mount *in situ* localization of *Tu-CPR* in *T. urticae* (i, ii, iv-vii, anti-sense probe). (i) adult female, (ii) adult male, (iii) adult female (hybridization with sense probe, background control). (iv-vii) enlarged views of (iv) epithelium, (v) ovaries and (vi and vii) digestive cells of adult female, taken from independently stained mites. **G**-**I**, Effects of dsRNA-*Tu-CPR*. **G**, Relative level of *Tu-CPR* transcript normalized with *RP49* in dsRNA-treated Col-a mites (mean ± SEM, *n* = 8). **H**, The activity of cytochromes P450 in dsRNA-treated Col-a mites upon feeding on Col-0 plants (mean ± SEM, *n* = 5). **I**, Fecundity of dsRNA-treated Col-a mites feeding on Col-0 or bean plants. Fecundity was measured over two days (3^rd^ and 4^th^ dpi) and data are presented as mean number of eggs laid by a female mite per day ± SEM, *n* = 30 for Col-0 plants and *n* = 12 for bean plants. See also Supplemental Figure S5. Asterisks indicate a significant difference between treated and control samples (unpaired Student’s t test: *P < 0.05, **P < 0.01, ***P < 0.001). Scale bars in **F**: (i-iii) 20 μm; (iv-vii) 100 μm. Individual sample values are shown as open circles for *n* ≤ 10.

The application of DEF reduced esterase activities by 32% but did not affect the fitness of Col-a mites when feeding on either Col-0 or *cyp79b2 cyp79b3* plants (Supplemental Figure S4), indicating that esterases may not be required for mite adaptation to *Arabidopsis*. The application of DEM decreased the GST enzymatic activity by 43% and had a minor but significant effect on the fecundity of Col-a mites (Supplemental Figure S4), suggesting that GST activity is required for high Col-a performance on *Arabidopsis*.

The application of PBO significantly reduced the P450 activity (25%) and dramatically reduced (44%) the fecundity of Col-a mites feeding on Col-a host plants, and to a lesser extent when they fed on *cyp79b2 cyp79b3* plants (Figure 5A,B). Since PBO does not interfere with all P450s equally and also inhibits esterases in some arthropods (Feyereisen, 2014), we applied another structurally unrelated P450 inhibitor, TCPPE, to confirm that the observed effects were due to the decreased P450 activity. TCPPE significantly inhibited P450 enzyme activity in Col-a mites (13%) and dramatically reduced the fecundity of Col-a mites fed on Col-0 (39%) and *cyp79b2 cyp79b3* plants (33%), (Figure 5C,D). In addition, the fitness of bean-a mites decreased by 13% when they fed on *cyp79b2 cyp79b3* TCPPE-treated plants (Figure 5D).

### P450 activity is required for TSSM adaptation to *Arabidopsis*

As P450 inhibitors were applied directly to *Arabidopsis* leaves, there is a possibility that PBO/TCPPE perturbed *Arabidopsis* defense physiology and only secondarily affected mite fecundity. For example, the synthesis of JA metabolites (Aubert *et al*., 2015; Heitz *et al*., 2012; Widemann *et al*., 2015) and Trp-derived secondary metabolites (Hull *et al*., 2000) occur via pathways that include many plant P450s. Therefore, to confirm the requirement of mite P450s in TSSM adaptation to *Arabidopsis*, we used recently developed RNAi gene silencing protocol in TSSM (Bensoussan *et al*., 2020; Suzuki *et al*., 2017) to silence the P450 pathway in mites.

Because the *CYP* (PF00067) gene family has over 100 *CYP* genes (Grbic *et al*., 2011; Mistry *et al*., 2020), it is difficult to characterize the involvement of individual P450s in mite adaptation to *Arabidopsis* using RNAi. To circumvent this hurdle, we instead silenced the expression of *NADPH-cytochrome P450 reductase* (*Tu-CPR*), a co-enzyme required for the catalytic reactions carried out by microsomal P450s (Masters and Okita, 1980). *Tu-CPR* (*tetur18g03390)* is a single copy gene (Grbic *et al*., 2011) constitutively expressed in all tissues of adult spider mites, including the midgut epithelial and digestive cells (Figure 5F) where digestion and detoxification of dietary xenobiotics are expected to take place (Bensoussan *et al*., 2018). Application of dsRNA-*Tu-CPR* to Col-a mites resulted in decreased expression of *Tu*-*CPR* (Figure 5G). Consistent with the Tu-CPR function as an essential component of the P450 enzyme complex, P450 activity was reduced by 38% in the *Tu-CPR* silenced mites relative to mites exposed to dsRNA-NC (Figure 5H). Strikingly, the silencing of *Tu-CPR* reduced the fecundity of Col-a mites by almost 50% when they fed on Col-0 plants (Figure 5I). Similar phenotypes were obtained upon the application of a second, independent, dsRNA-*Tu-CPR-1* fragment (Supplemental Figure S5), confirming the specificity of the observed phenotypes to a loss of *Tu-CPR* function and the requirement of P450 activity for the adaptation of Col-a mites to *Arabidopsis*. The reduced P450 activity, through the application of P450 inhibitors or dsRNA-*Tu-CPR*, did not affect TSSM fecundity when mites fed on bean plants (Figure 5E,I and Supplemental Figure S5), demonstrating that P450 activity is specifically required for mites’ ability to use *Arabidopsis* as a host.

## Discussion

The innate ability to perceive and respond to plant defenses in a process of physiological acclimation is assumed to enable broad-generalist herbivores to utilize a wide range of host plants. On the other hand, genetic adaptation is presumed to enable specialist herbivores to evolve novel adaptation traits that efficiently counteract the defenses of a narrow range of host plants (Barrett and Heil, 2012; Despres *et al*., 2007; Heidel-Fischer and Vogel, 2015). However, our understanding of the host-shift responses in composite generalist herbivores that exist as complexes of host-specialized populations is not well understood. *Tetranychus urticae* is an example of a composite generalist herbivore that feeds on over thousand plant hosts. Individual mite populations thrive on a subset of potential hosts but can rapidly adapt to new plant hosts (Agrawal *et al*., 2002; *Fellous et al., 2014*; Fry, 1989; Gould, 1979; Magalhaes *et al*., 2007; Wybouw *et al*., 2015). Thus, mite host adaptability is one of the key features underlying *T. urticae* extremely wide host range.

*T. urticae*, like other polyphagous herbivores, acclimates to host shift by reprogramming its salivary and detoxification complements (Figure 4 and (*Dermauw et al., 2013*; Jonckheere *et al*., 2018; Jonckheere *et al*., 2016; Villarroel *et al*., 2016; Wybouw *et al*., 2015; Zhurov *et al*., 2014)). However, exposure to initially unfavorable host plants over several generations enables *T. urticae* to dramatically increase its performance (Figure 2 and (Fry, 1989; Gould, 1979; Magalhaes *et al*., 2007; *Wybouw et al., 2015*). Here, we used *Arabidopsis* and bean-adapted (bean-a) mites to investigate mechanisms that mediate the formation of host-adapted populations in this generalist herbivore. *Arabidopsis* is a non-preferred host for the bean-a mite population, Figure 1. However, under experimental selection, bean-a mites gave rise to *Arabidopsis*-adapted *cyp*-a and Col-a populations, Figure 2. Matching the complexity of *Arabidopsis* defenses, *Arabidopsis*-adapted populations evolved adaptation traits capable of counteracting *CYP79B2 CYP79B3*-independent *Arabidopsis* defenses (present in both *cyp*-a and Col-a mites) and Trp-derived defenses (present only in Col-a mites). Remarkably, these complex adaptation traits were established within 25 generations of mite *Arabidopsis* selection.

Suppression of plant defenses has been proposed to be one of the mechanisms of mite host adaptation (Blaazer *et al*., 2018; Jonckheere *et al*., 2018; Jonckheere *et al*., 2016; Villarroel *et al*., 2016). Attenuation of host induced responses has been described for several spider mites species in their interactions with the *Solanum lycopersicum* (tomato) plant host (Glas *et al*., 2014; Kant *et al*., 2008; P. Godinho *et al*., 2016; Sarmento *et al*., 2011). The prevalence of the ability to suppress plant responses across tomato-adapted *T. urticae* populations is currently not known, nor is its presence in other *T. urticae* host-adapted populations. Even though the interference with either JA-biosynthesis or its signalling could have been an effective way to attenuate a whole range of *Arabidopsis* defense compounds, (Figure 1, (Widemann *et al*., 2021; *Zhurov et al., 2014*)), our data suggest that *Arabidopsis*-adapted mites did not interfere with JA-regulated *Arabidopsis* defenses. *Arabidopsis*-adapted mites, just like the ancestral bean-a population, had greater fitness on *cyp79b2 cyp79b3* than on Col-0 plants (Figure 2). They also induced the expression of JA-responsive genes and accumulated I3M to levels similar to those induced by the bean-a mites (Figure 3). This is surprising, as ancestral bean-a strain used in this study previously gave rise to tomato-adapted populations that, in the process, gained the ability to suppress tomato mite-induced responses (Wybouw *et al*., 2015), raising the question if and how host defenses affect the mechanism of TSSM adaptation. Instead, we provided multiple independent lines of evidence indicating that *Arabidopsis*-adapted mites evolved metabolic resistances to counteract *Arabidopsis* defenses.

The pharmacological experimental treatments identified the requirement of GSTs for the fitness advantage of Col-a over bean-a mites when they fed on Col-0 plants (Supplemental Figure S4). GST-based detoxification has been associated with the ability of numerous generalist herbivores to counteract the anti-herbivore effects of glucosinolates, considered to be the major class of *Arabidopsis* defense compounds (Gloss *et al*., 2014; Jeschke *et al*., 2016; Schramm *et al*., 2012; Wadleigh and Yu, 1988). Aliphatic and indole glucosinolates are the most abundant classes of these compounds in *Arabidopsis* (Brown *et al*., 2003). Herbivores have differential susceptibility to aliphatic and indole glucosinolates, so that some are exclusively affected by the aliphatic (Müller *et al*., 2010), some by the indole (Kim and Jander, 2007), and some by both classes of glucosinolates (Jeschke *et al*., 2017; Müller *et al*., 2010). Aliphatic glucosinolates are ineffective in restricting mite herbivory (Zhurov *et al*., 2014). Instead, the antifeedant effects of indole glucosinolates on *T. urticae* have been recently characterized (Widemann *et al*., 2021), raising a possibility that *Arabidopsis* defensive indole glucosinolate breakdown products may be detoxified by mite adaptation-associated GST(s).

The pharmacological and RNAi reverse genetics experiments pointed to the critical requirement of P450 activities for the adaptation of Col-a mites to *Arabidopsis* (Figure 5, Supplemental Figure 4). We identified two distinct modes of P450 activities. One corresponds to the initial mite response to the shift from bean to *Arabidopsis* that is similar in both ancestral and *Arabidopsis*-adapted mites (Figure 4). The inhibition of these P450s had a limited but significant effect on the fecundity of bean-a ancestral TSSM population when these mites were challenged on the *Arabidopsis* host (Figure 5D). As bean-a mites had no prior exposure to *Arabidopsis* plants, the effect of P450 inhibitors on their fecundity uncovers the limited contribution of initially induced P450s to mite fitness on *Arabidopsis*. Since *Arabidopsis*-adapted mites retained responsiveness to shift from bean to *Arabidopsis* plants, Figure 4, it is expected that these P450s enable limited mite adaptation to *Arabidopsis* defenses ((Zhurov et al., 2014) and Figure 5). This is consistent with the physiological acclimation described for broad-generalists whereby exposure to plant xenobiotics is assumed to result in the transcriptional induction of genes encoding enzymes that recognize a wide range of substrates and have low enzymatic catalytic activity (Halon *et al*., 2015; Li *et al*., 2004; Mathers *et al*., 2017; Sasabe *et al*., 2004; Shi *et al*., 2018; Snoeck *et al*., 2019; Wang *et al*., 2018; Wen *et al*., 2006). The other more significant contribution of P450-mediated metabolic resistance is realized through the action of highly effective P450 detoxification responses against *Arabidopsis* defenses that are exclusively associated with mite adaptation to *Arabidopsis*. The reduction of P450 activity in Col-a mites diminished their fitness advantage over ancestral bean-a mites, demonstrating that adaptation-associated P450 activity is the main contributor to mite resistance to *Arabidopsis* defenses (Figure 5B,D,I and Supplemental Figure S5). The inhibition of P450 activity reduced Col-a mite fecundity when they fed on both Col-0 and *cyp79b2 cyp79b3* plants (Figures 5B,D), suggesting that adaptation-associated P450s mainly counteract CYP79B2 CYP79B3-independent *Arabidopsis* defenses whose identity is currently not known. However, the requirement of P450 activity for the adaptation of green peach aphid *M. persicae* to *Arabidopsis* indole glucosinolates (Ji *et al*., 2021) raises the possibility that mite adaptation-associated P450s may also counteract the toxicity of indole glucosinolate-associated defensive compounds.

In summary, as demonstrated here for the *Arabidopsis*-adapted mite populations, *T. urticae* can evolve resistance against complex plant defenses within a mere 25 generations. *Arabidopsis*-adapted mites retained the ability to acclimate to shift from bean to *Arabidopsis*. Such transcriptional plasticity, characteristic of broad-generalists, is shared between ancestral and *Arabidopsis*-adapted mites and has a small contribution to mite fitness on *Arabidopsis*. At present, it is not clear if these initial responses are common stress/xenobiotic responses to any novel host, nor if they are required for mite host adaptation. If they are, they may provide initial fitness benefit upon mites’ shift to a new host that may facilitate the evolution of the host adaptation traits. We further identified the highly effective P450 detoxification responses that are required for mite adaptation to *Arabidopsis*. This pattern of metabolic counteraction is reminiscent of strategies deployed by host-specialised herbivores, in which host adaptation is associated with either nucleotide substitutions that enhance the enzyme activity (Li *et al*., 2007; Schuler, 2011) or with the overexpression of detoxification genes (Bass *et al*., 2013). The *CYP* gene family underwent a lineage-specific expansion in TSSM and consists of 115 genes in bean-a mites (Grbic *et al*., 2011; Mistry *et al*., 2020). Intraspecific genetic variability is expected at *CYP* loci as some appear as clusters of duplicated genes that are prone to the formation of chimeric genes or copy number variation. Comparative genome and transcriptome analysis of *Arabidopsis*-adapted and ancestral TSSM populations may identify candidate adaptation-required *CYP* gene(s). In humans, for example, only 15 out of 57 P450s are involved in xenobiotic metabolism (Handschin and Meyer, 2003). The comparative sequence and functional analysis of candidate *CYPs* will reveal if adaptation-required P450 activity is achieved by altering gene regulation, copy number, or through substitution(s) that increase enzyme activity and if these changes resulted from the selection within the genetic pool of founding population, through *de novo* mutations or epigenetic changes.

## Materials and Methods

### Plant growing and mite rearing conditions

Columbia-0 (Col-0) seeds were obtained from the Arabidopsis Biological Resource Center (ABRC, Ohio State University), *cyp79b2 cyp79b3* (Zhao *et al*., 2002) from B. A. Halkier (University of Copenhagen, Denmark), *myc2 myc3 myc4* (Fernandez-Calvo *et al*., 2011), from R. Solano (Universidad Autónoma de Madrid) and bean (*Phaseolus vulgaris*, cultivar California Red Kidney) from Stokes, Thorold, Ontario, Canada. Plants were grown under 100 to 150 μmol m^−2^ sec^−1^ cool-white fluorescent light at 24°C with a 10-h/14-h (light/dark) photoperiod in controlled growth chambers. *Tetranychus urticae*, London reference strain (Grbic *et al*., 2011), were reared on bean plants at 24°C, 60% relative humidity, and with a 16-h/8-h (light/dark) photoperiod for >10 years.

### Experimental evolution

Approximately 1,000 randomly chosen adult bean-a female mites were transferred to 3-week-old *cyp79b2 cyp79b3* or Col-0 *Arabidopsis* plants that were replaced biweekly. Mite populations were allowed to propagate for 18 months, generating six selected lines: #1-#3 independent lines each selected on *cyp79b2 cyp79b3* and Col-0 plants, referred to as *cyp*-a and Col-a lines respectively. Selected lines were reared on their corresponding plant hosts under the same conditions as the ancestral strain. *cyp*-a (#3) and Col-a (#3) were used for the follow-up experiments.

### Mite performance analysis

Twenty 3-day-old adult female mites were transferred to each of the experimental plants (bean, *cyp79b2 cyp79b3* and Col-0) and total population size at seven days post-inoculation was evaluated in direct (mites were moved directly from their rearing hosts to experimental plants) and indirect (mites were reared on bean plants for two generations before their transfer to the experimental plants) transfer regimes. The experiment was performed in four biological replicates for each selected line. Differences between selected line population sizes were determined using a one-way ANOVA using mite strain as the main effect, followed by Tukey’s honestly significant difference (HSD) test. All other experiments were performed with mites that were propagated for two generations on bean plants, thus, in indirect transfer regime.

### Plant damage analysis

Leaf damage of *cyp79b2 cyp79b3* and Col-0 *Arabidopsis* plants upon feeding of bean-a, *cyp*-a, and Col-a mite strains was performed as previously described (Zhurov *et al*., 2014). Briefly, ten adult female mites were placed on the rosette of *Arabidopsis* plants and allowed to feed for 3 days before plants were cut at the base of the rosette. The adaxial side of the rosette was scanned using a Canon CanoScan 8600F model scanner (Canon U.S.A. Inc., Melville, NY, U.S.A) at a resolution of 1200 dpi and a brightness setting of +25. Scanned plants were saved as .jpg files for subsequent analysis in Adobe Photoshop 5 (Adobe Systems, San Jose, CA). The experiment was performed using six biological replicates/trial, and in three experimental trials, *n* = 18. Differences in plant damage between mite strains on different hosts were determined using a three-way ANOVA, using mite strain, plant host, and experimental trial as main effects, including interaction terms, followed by Tukey’s HSD test. The inclusion of experimental trial and its potential interactions with main effects was used both to control for the effect of trial and to test for reproducibility across trials, as suggested by (Brady *et al*., 2015). This statistical approach was used in all following analyses.

### Mite fecundity assay

Fecundity over two days of bean-a mites on Col-0, *cyp79b2 cyp79b3*, and *myc2 myc3 myc4*, and fecundity over six days of bean-a, *cyp*-a, and Col-a mites on *cyp79b2 cyp79b3* or Col-0 *Arabidopsis* plants were assessed as previously described (Widemann *et al*., 2021). The experiment included five to ten biological replicates for each treatment that were repeated in three experimental trials. A two- and three-way ANOVA was performed respectively, using the mite strain, plant genotype, and experimental trial as main effects including relevant two-way interaction terms when required. For the mite fecundity assay on MeJA-treated Col-0 leaves, the leaves from five-week-old Col-0 plants were sprayed 6 times over 48 h with methyl jasmonate (MeJA) (Sigma-Aldrich, Cat # 392707, 500 μM in ethanol 0.4 % (v/v)), as previously described (Feldberg *et al*., 2009). On the second day of treatment, fully-elongated leaves were detached and a day later infested with 10 adult female bean-a mites whose fecundity was recorded after 24 h. The experiment included nine biological replicates and was repeated in two experimental trials. Differences in fecundity were detected by a two-way ANOVA, using the treatment and experimental trial as main effects including an interaction term.

### Real-Time Quantitative Reverse Transcription-PCR and metabolic analyses

For plant marker gene and metabolite analyses, ten three-day-old adult female bean-a, *cyp*-a, and Col-a mites, propagated for two generations on bean plants, were allowed to feed on Col-0 plants for 24 h after which whole rosettes were collected. For TSSM marker gene analysis, samples of 100 three-day-old adult female bean-a, *cyp*-a, and Col-a mites, initially propagated for two generations on bean plants and then transferred to bean, *cyp79b2 cyp79b3*, or Col-0 plants for 24 h, were collected. Both experiments were replicated in three biological replicates/trial and three independent trials. Preparation of RNA, cDNA, and the real-time quantitative reverse transcription-PCR analysis was performed as previously described (Zhurov *et al*., 2014). Primer sequences and amplification efficiencies (E) used in qPCR are shown in Supplemental Table 1. *PEROXIN4* (*AT5G25760*) and *RP49* (*tetur18g03590*) were used as the reference genes for *Arabidopsis* and mite genes, respectively. Ct values of three technical replicates were averaged to generate a biological replicate Ct value. Plotting and statistical analysis were performed as described previously (Rieu and Powers, 2009). The quantification of plant metabolites (JA, JA-Ile, and I3M) was performed as described in Zhurov et al., 2014. Briefly, jasmonates (JA and JA-Ile) were analyzed in aqueous extracts by mass spectrometry monitoring their specific precursor-to-product ion transition in a triple quadrupole mass spectrometer (Micromass Ltd. UK). Quantitation was achieved after external calibration with standards of known amount considering the analyte/internal standard ratio (dihydrojasmonic acid, added before extraction). Moreover, 3-indolylmethyl glucosinolate (I3M) was identified and analyzed following a non-targeted approach using a hybrid quadrupole-time-of-flight (QqTOF) mass spectrometer. Identification of I3M was performed after inspection of its specific mass spectrum in negative electrospray ionization mode: 447.053 [M-H]^−^, 253.03 [C_10_H_9_N_2_O_4_S]^−^, 96.95 [HSO_4_]^−^. Relative quantitation of I3M was attained by dividing peak areas of I3M and internal standard (biochanin A, 283.07 [M-H]^−^). The resulting value was again divided by the actual sample weight expressed in mg. Mite marker gene data were analyzed by a two-way ANOVA with mite strain and plant host as main effects including an interaction term, followed by Tukey’s HSD test. For plant marker gene and metabolite analysis, the data were analyzed by a two-way ANOVA, using mite strain and experimental trial as main effects including an interaction term, followed by Tukey’s HSD test. For a graphical representation of the expression of plant marker genes, the NRQ data were further normalized to the ‘no mite’ control sample, which was therefore set to one. Thus, these results represent fold change differences in the expression of marker genes relative to ‘no mite’ control.

### Determination of cytochrome P450 monooxygenase (P450), esterase and glutathione-S-transferase (GST) activity

To perform enzymatic activity assays, 3-day-old spider mite females of each Col-a, *cyp*-a, and bean-a strains, reared for two generations on bean plants, were placed on bean, *cyp79b2 cyp79b3*, or Col-0 plants. After 24 h, 200 females from each treatment were collected and were used for protein extraction and enzymatic assays. Mites were homogenized in 700 μL of 100 mM phosphate buffer, pH 7.6. The concentration of total protein mite extracts was determined using the Quick Start Bradford Protein Assay (Quick start Bradford 1x dye reagent, Bio-Rad, Cat# 500-0205), with Bovine Serum Albumin (Sigma-Aldrich, Cat # A7906) as the standard. Twenty, 10, and 10 μg of protein per reaction were used for measuring the enzymatic activity of glutathione-S-transferases (GSTs), P450 monooxygenases, and esterases, respectively. The enzymatic activities were assessed spectrophotometrically using 1-chloro-2,4-dinitrobenzene (CDNB) (for GSTs) (Habig and Jakoby, 1981), 7-ethoxy-4-trifluoromethylcoumarin (7-EFC) (for P450s) (Buters *et al*., 1993), and 4-nitrophenyl acetate (pNPA) (for esterases) (Park *et al*., 1961), as substrates. For GST and esterases, the linearity of the reaction was determined by plotting the absorbance values against time over 5 minutes period and calculation of samples activity was performed in the linear range. All assays were corrected for the occurrence of non-enzymatic substrate transformation in blank samples in which the protein solution was replaced by the buffer. Enzymatic activities were determined in four independent samples with three technical replicates per sample. Differences in enzymatic activity between mite strains on different plant hosts were detected using a two-way ANOVA, with mite strain and plant host as main effects including an interaction term, followed by Tukey’s HSD test.

### Application of enzyme inhibitors

Solutions of 2000 mg/L diethyl maleate (DEM, an inhibitor of glutathione S-transferase (GST) activity), 100 mg/L S,S,S tributyl-phosphorotrithioate (DEF, an inhibitor of esterase activity), 1000 mg/L piperonyl butoxide (PBO) and 1500 mg/L trichlorophenylpropynyl ether (TCPPE) (inhibitors of P450 activities) were used as they cause <10% mortality of bean-a mites (Supplemental Figure S3). Detached leaves of Col-0 *Arabidopsis* plants were dipped in one of the DEF, DEM, PBO or TCPPE solutions or water and acetone 0.1 % (v/v) as control. Upon drying, leaves were infested with either one adult bean-a or Col-a female mite and the number of eggs laid by each female per day was recorded for six days, or with ten 2-day-old adult Col-a female mites that were collected after 24 h. A pool of 100 spider mites was used to measure GST, esterase, and P450 activities in DEM-, DEF-, PBO- and TCPPE-treated spider mites, respectively. Fecundity was determined in three independent experimental trials with ten replicates per trial. Enzymatic activities were determined in five independent samples with three technical replicates per sample. Differences between means of fecundity and enzymatic activities of control and inhibitor-treated samples were calculated using unpaired Student’s t-tests.

### *In situ* hybridization

DIG-labelled probes were produced and the whole-mount *in situ* hybridization was performed according to previously published methods (Dearden *et al*., 2002). Images were collected using a Zeiss AxioCam HRc 412-312 camera mounted on a Zeiss Axioplan II microscope.

### RNAi of *Tu-CPR*

Two non-overlapping fragments (*Tu-CPR*, 645 nt) and (*Tu-CPR-F1*, 564 nt), complementary to the coding region of *Tu-CPR (tetur18g03390*) and dsRNA complementary to a 382 bp non-transcribed intergenic fragment (1690614-1690995 of the genomic scaffold 12) used as a negative control dsRNA (referred to as NC) were synthesized using primers listed in Supplementary Table 1 as described in (Suzuki *et al*., 2017). A BLAST search against the *T. urticae* genome confirmed that dsRNA sequences are unique. dsRNA solutions at concentrations of 500 ng/μL were supplemented with 6% (v/v) blue dye erioglaucine (McCormick, Sparks Glencoe, MD). Newly molted adult female mites were soaked in dsRNA/dye solutions at 20°C for 24 h (Suzuki *et al*., 2017). Post soaking, mites with visible blue dye in the posterior midgut were selected and were transferred in batches of 10 to either detached Col-0 leaves or bean leaf disks. Fecundity was determined in three independent experimental trials with ten replicates/trial as a number of eggs deposited by individual female mite on the 3^rd^ and 4^th^ day post soaking. For the analysis of the *CPR* expression, 100 adult female mites were treated with either dsRNA-*Tu-CPR*, dsRNA-*Tu-CPR-1*, or dsRNA-NC. Mites were allowed to feed on Col-0 leaves for four days. Thereafter, mites were selected based on visual phenotype (spotless) and were used for the analysis of *CPR* expression levels. The RT-qPCR was performed in eight (for dsRNA-*CPR*) and three (for dsRNA-*CPR-1*) experimental replicates, as described above. For the analysis of P450 activity in *CPR* silenced mites, 100 adult female mites were treated with either dsRNA-*Tu-CPR*, dsRNA-*Tu-CPR-1*, or dsRNA-NC. After four days of feeding on Col-0 leaves, mites were collected and the P450 activity was determined in five independent samples with three technical replicates per sample, as described above. Differences in fecundity between dsRNA-treated mites were detected using a two-way ANOVA, with mite treatment and experimental trial as main effects including an interaction term, followed by Tukey’s HSD test. Enzyme activity and RT-qPCR expression levels were compared between dsRNA-treated mites using unpaired Student’s t-tests. Independent mite samples were used for the fecundity assay and the analysis of transcript levels and P450 activity.

## Acknowledgments

The authors would like to thank Drs. Thomas Van Leeuwen and Rene Feyereisen for their constructive comments on pharmacological experiments, and, Pierre Hilson and Mark Bernards for critical comments on the manuscript.

## Supplemental Data

**Supplemental Table 1**. Gene-specific primer sequences used for real-time quantitative RT-PCR.

**Supplemental Figure S1.**
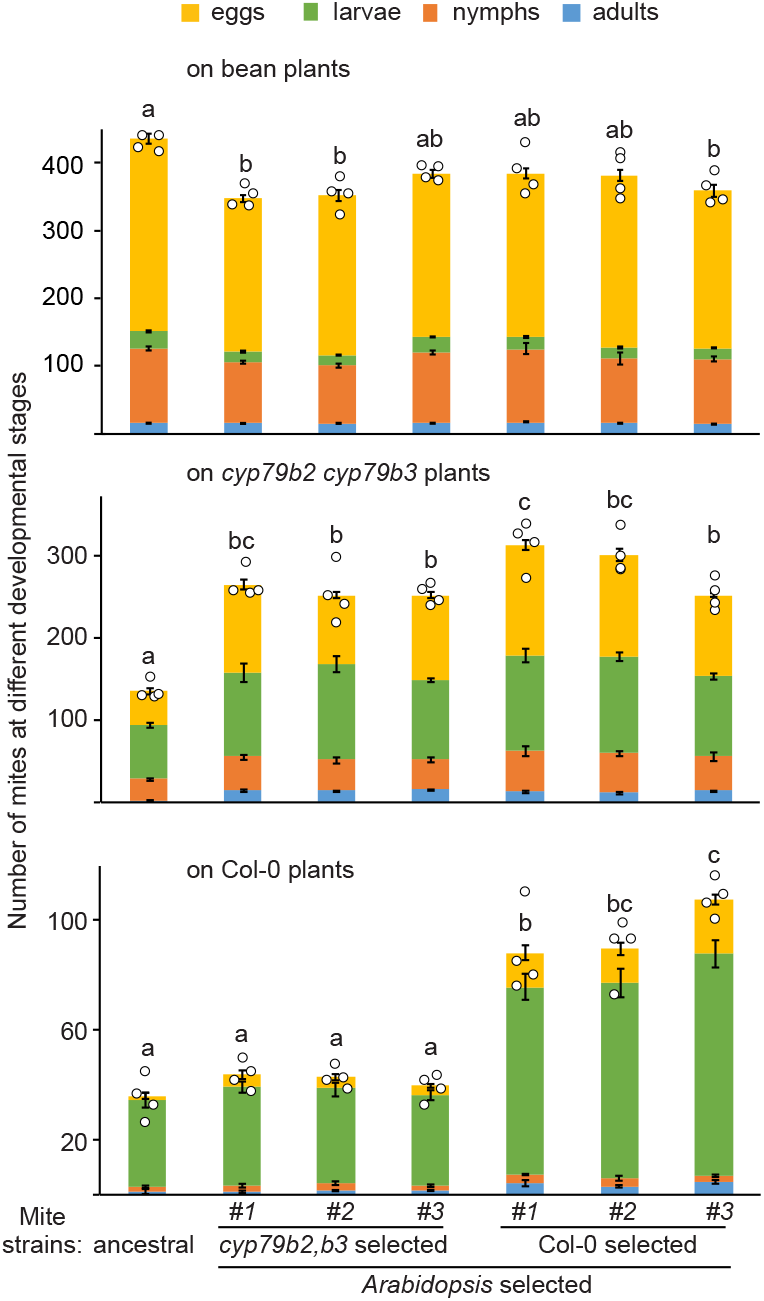
Experimental evolution of *Tetranychus urticae* adaptation to *Arabidopsis*. Performance of ancestral and selected populations on bean (top), *cyp79b2 cyp79b3* (middle) and Col-0 (bottom) plants. Mites were transferred directly from their respective rearing hosts to experimental bean, *cyp79b2 cyp79b3* and Col-0 plants. The performance was measured as the size of total population derived from twenty adult female mites and seven days post infestation. Data are represented as mean ± SEM, *n* = 4. Statistics were performed on total population counts. Different letters represent significant difference between means (Tukey’s HSD test, α = 0.05). Individual sample values are shown as open circles for n ≤ 10.

**Supplemental Figure S2.**
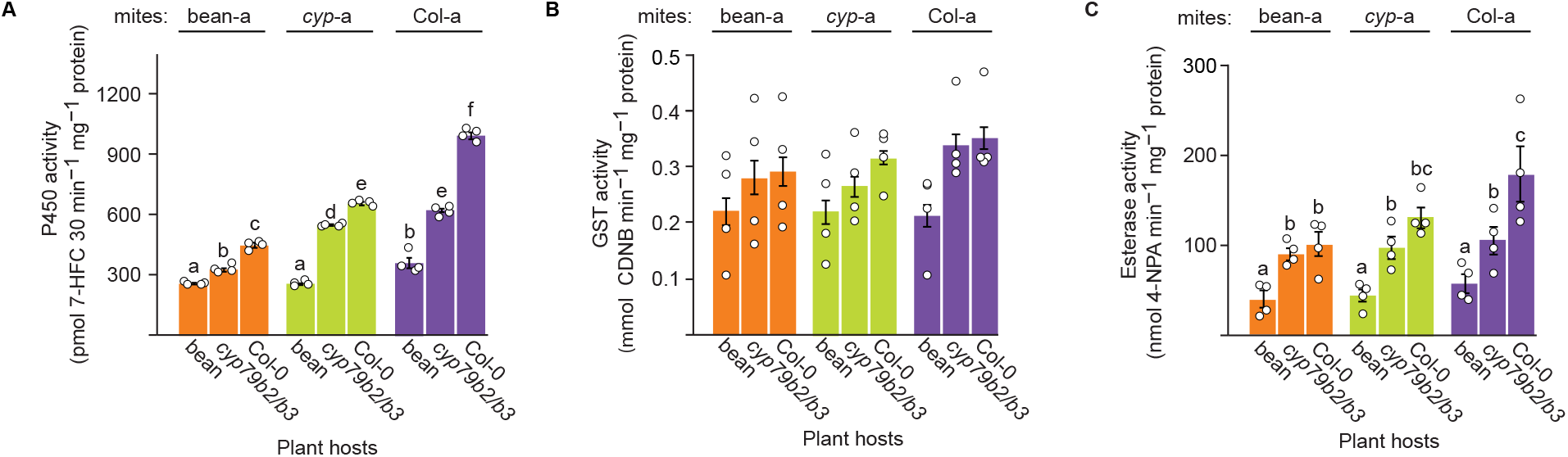
The activities of cytochromes P450 (**A**), glutathione-S-transferases (**B**) and esterase (**C**) in bean-a, *cyp*-a and Col-a mites (reared on beans for two generations) feeding on bean, *cyp79b2 cyp79b3* or Col-0 plants. Data are represented as mean ± SEM, *n* = 4. Different letters represent significant differences between means (Tukey’s HSD test, α = 0.05), following three-way ANOVA. Plant host - mite strain interaction was significant for the activity of cytochromes P450 (F = 103.93, P = 5.451e-16). Individual sample values are shown as open circles for n ≤ 10.

**Supplemental Figure S3.**
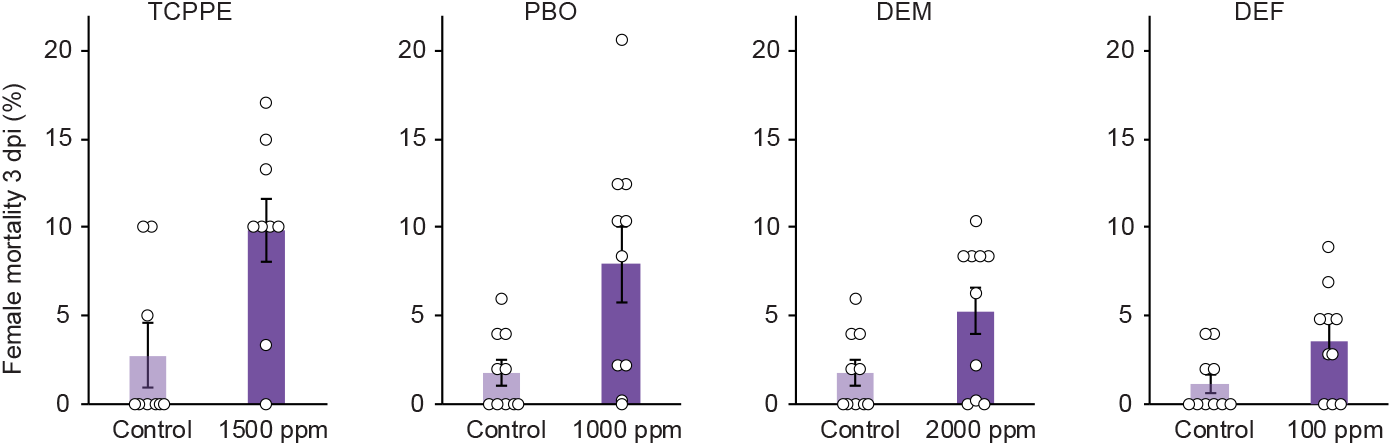
Selection of experimental concentration of enzyme inhibitors. Data are presented as mean mortality of a female mite at 3 dpi ± SEM (for treatment and control *n* = 10 for PBO, DEM and DEF experiments, and *n* = 9 for TCPPE experiment). Individual sample values are shown as open circles for n ≤ 10.

**Supplemental Figure S4.**
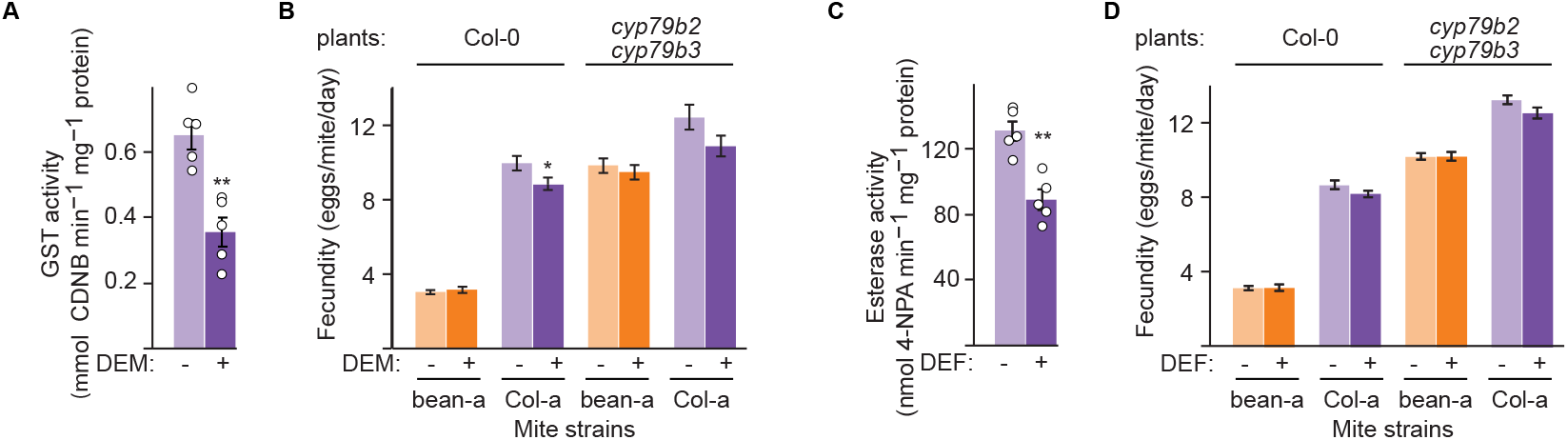
The requirements of esterase and glutathione-S-transferase (GST) activities for *Tetranychus urticae* adaptation to *Arabidopsis*. The GST (**A**) and esterase (**C**) activities in Col-a mites feeding on Col-0 plants after the application of diethyl maleate (DEM, an inhibitor of GST activity, in **A**) and S,S,S tributyl-phosphorotrithioate (DEF, an inhibitor of esterase activity, in **C**). Data are represented as mean ± SEM, *n* = 5. **B** and **D**, Effects of DEM (in **B**) and DEF (in **D**) treatments on fecundity of bean-a and Col-a mites upon feeding on Col-0 and *cyp79b2 cyp79b3* plants. Fecundity was assessed as mean number of eggs laid by a female mite in six days ± SEM, *n* = 30. Asterisks in panels **A**-**C** indicate a significant difference between treated and control samples (unpaired Student’s t test: *P < 0.05, **P < 0.01). Individual sample values are shown as open circles for n ≤ 10.

**Supplemental Figure S5.**
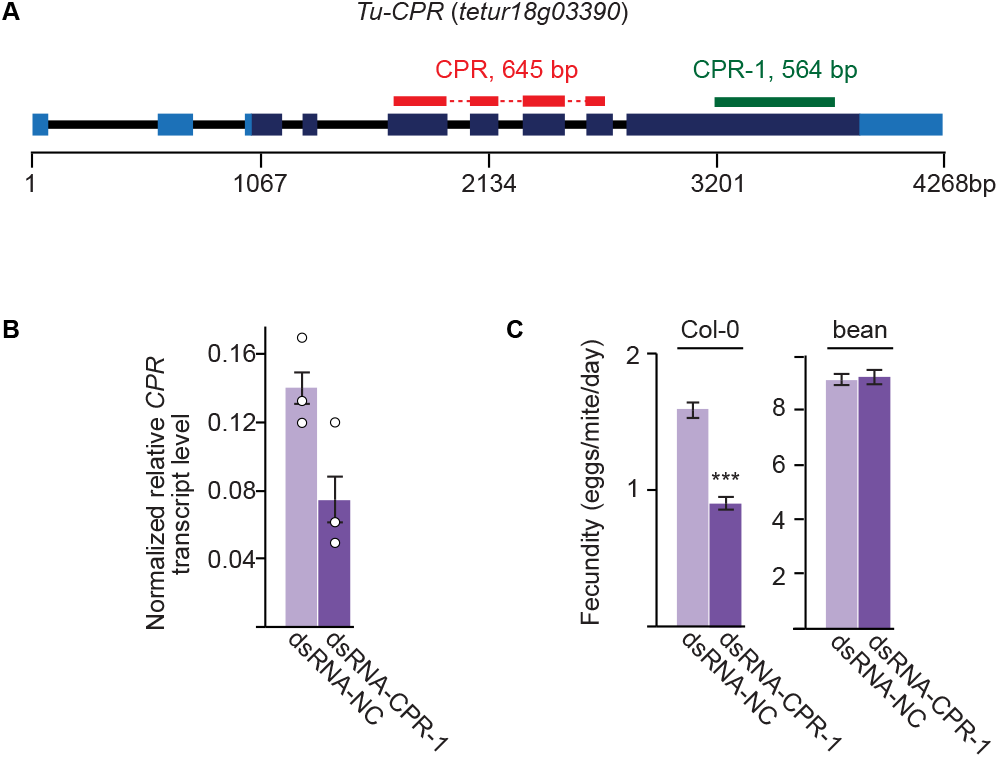
RNAi silencing of *Tu-CPR* gene. **A**, A schematic of the *Tu-CPR* locus. DNA sequences used for the generation of dsRNA-*Tu-CPR* are shown in red (fragment CPR, 645 bp), and green (fragment CPR-1, 564 bp). UTR and coding sequences are shown as light and dark blue boxes, respectively. **B**, The effect of dsRNA-*Tu-CPR*. Relative expression of *Tu-CPR* normalized with *RP49* in dsRNA treated Col-a mites (mean ± SEM, *n* = 8, Student’s t test, ns). **C**, Fecundity of dsRNA treated Col-a mites feeding on Col-0 and bean plants. Fecundity was measured over two days (3 and 4 dpi) and data are presented as mean number of eggs laid by a female mite per day ± SEM, *n* = 30 for Col-0 plants and *n* = 12 for bean plants. Data are represented as mean ± SEM (Student’s t test *** P < 0.001). Individual sample values are shown as open circles for n ≤ 10.

**Supplemental Table 1.**
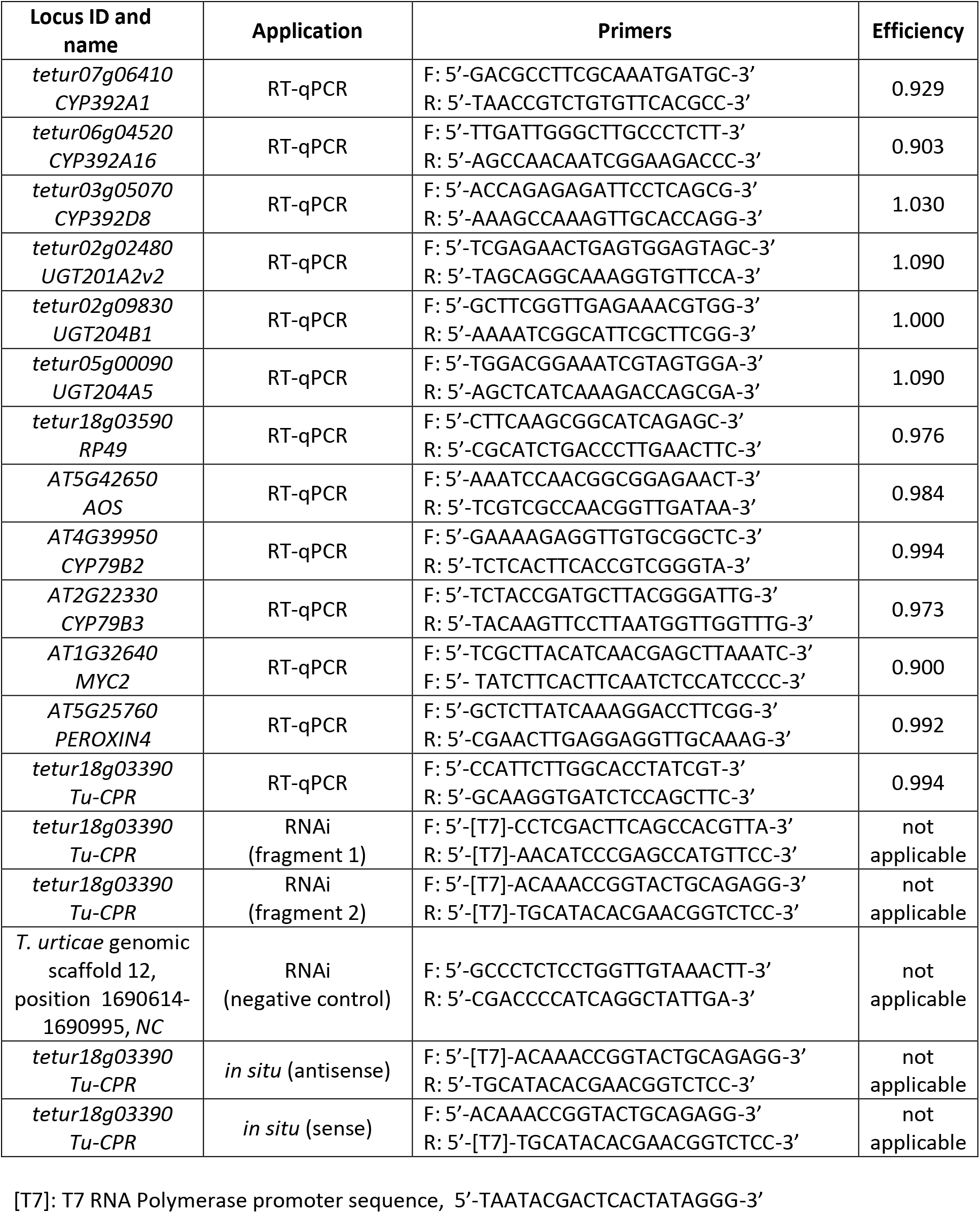
Gene-specific primer sequences used for the analysis of levels of gene expression (RT-qPCR), synthesis of dsRNA (RNAi) and preparation of probe for the *in situ* transcript localization.

